# A Computational Model of Bidirectional Axonal Growth in Micro-Tissue Engineered Neuronal Networks (micro-TENNs)

**DOI:** 10.1101/369843

**Authors:** Toma Marinov, Liang Yuchi, Dayo O. Adewole, D. Kacy Cullen, Reuben H. Kraft

## Abstract

Micro-Tissue Engineered Neural Networks (Micro-TENNs) are living three-dimensional constructs designed to replicate the neuroanatomy of white matter pathways in the brain, and are being developed as implantable microtissue for axon tract reconstruction or as anatomically-relevant *in vitro* experimental platforms. Micro-TENNs are composed of discrete neuronal aggregates connected by bundles of long-projecting axonal tracts within miniature tubular hydrogels. In order to help design and optimize micro-TENN performance, we have created a new computational model including geometric and functional properties. The model is built upon the three-dimensional diffusion equation and incorporates large-scale uni- and bi-directional growth that simulates realistic neuron morphologies. The model captures unique features of 3D axonal tract development that are not apparent in planar outgrowth, and may be insightful for how white matter pathways form during brain development. The processes of axonal outgrowth, branching, turning and aggregation/bundling from each neuron are described through functions built on concentration equations and growth time distributed across the growth segments. Once developed we conducted multiple parametric studies to explore the applicability of the method and conducted preliminary validation via comparisons to experimentally grown micro-TENNs for a range of growth conditions. Using this framework, this model can be applied to study micro-TENN growth processes and functional characteristics using spiking network or compartmental network modeling. This model may be applied to improve our understanding of axonal tract development and functionality, as well as to optimize the fabrication of implantable tissue engineered brain pathways for nervous system reconstruction and/or modulation.

## Introduction

Various neural tissue engineering tools have been created to model and study the development of neuronal networks *in vitro*. Among them are micro-tissue engineered neural networks (micro-TENNs), which are three-dimensional (3D) living constructs comprised of long-projecting axonal tracts and discrete neuronal populations within a microscopic, hollow hydrogel cylinder (microcolumn) filled with an extracellular matrix (ECM) [1]. Preformed clusters of neuronal cell bodies (aggregates) are housed at one or both ends of the microcolumn, with axons growing longitudinally through the hydrogel lumen (**Figure 1**). This segregation of long axonal tracts and neuronal cell bodies approximates the network architecture of the central nervous system by replicating the anatomy of gray matter and white matter pathways referred to as the “connectome”. Micro-TENNs may be fabricated with a range of neuronal subtypes and physical properties, yielding a controllable yet biofidelic microenvironment for studying 3D neural networks *in vitro*. As such, micro-TENNs are being developed in parallel as (1) self-contained, bioengineered implants to reconstruct compromised pathways in the brain, and (2) biofidelic test-beds for studying various neuronal phenomena (e.g. growth, synaptic integration, circuit development, pathological responses) [1]–[5]. Towards the former, prior work has shown that micro-TENNs are capable of survival, maintenance of architecture, neurite outgrowth, and host/implant synaptic integration out to at least 1 month following implant in adult rats [3]–[6].

**Figure 1.**
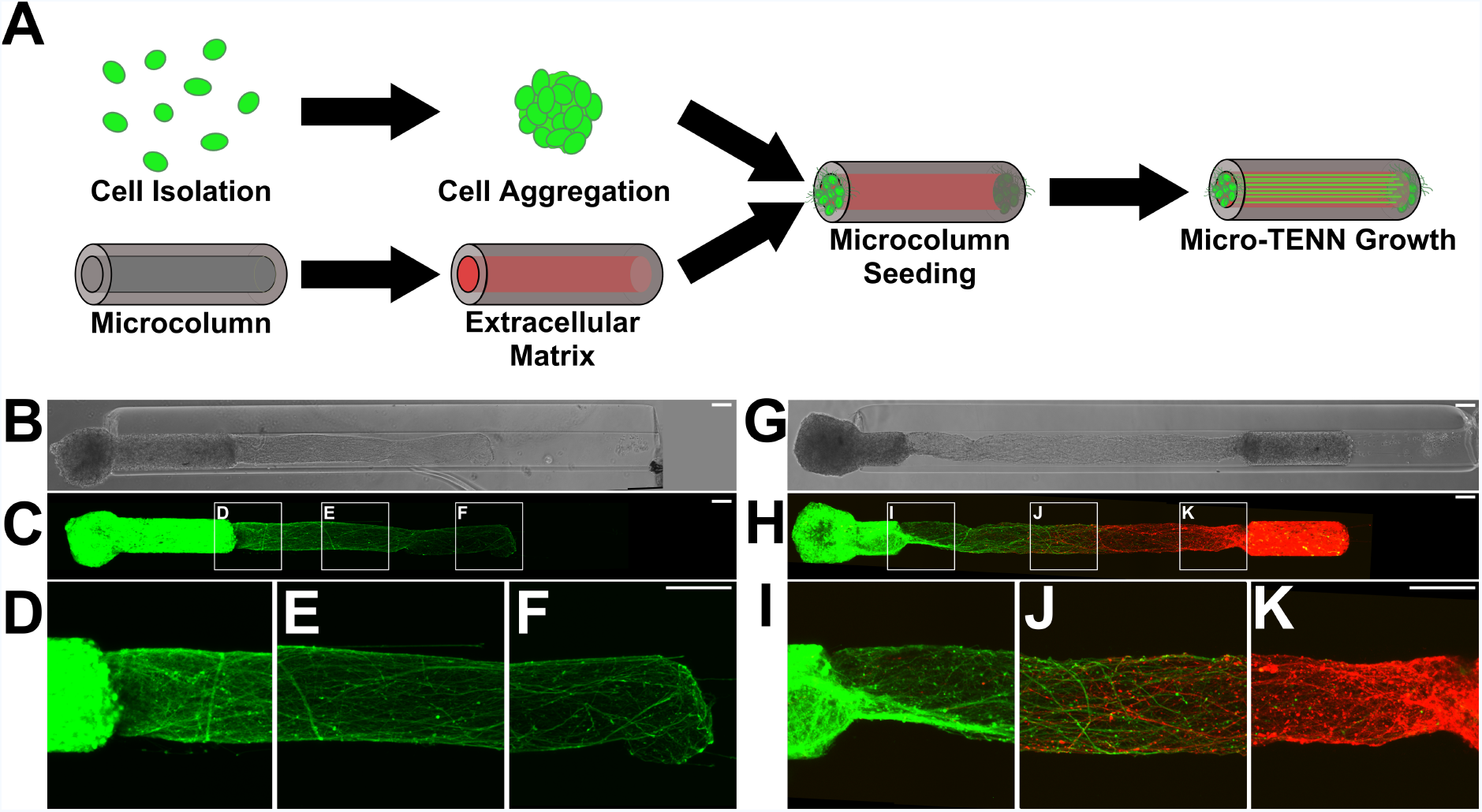
Micro-TENNs as living, 3D neuronal-axonal constructs. **(A)** Micro-TENNs are fabricated in a three-step process. Neurons are isolated and forced into spherical aggregates via gentle centrifugation (top). Simultaneously, single-channel microcolumns are cast from hydrogel to a predetermined inner and outer diameter and filled with an extracellular matrix (ECM) comprised of collagen and laminin (bottom). Next, ECM-filled microcolumns are seeded with either 1 aggregate or 2 aggregates to form either unidirectional or bidirectional micro-TENNs, respectively. Micro-TENNs are then grown *in vitro*. **(B)** Phase microscopy image of a unidirectional, GFP-positive micro-TENN at 5 days in vitro (DIV). **(C-F)** Confocal micrograph of same micro-TENN from (B). Axons can be seen projecting from the neuronal aggregate **(D)** and extending through the ECM-filled microcolumn **(E, F)**. **(G)** Phase micrograph of a bidirectional micro-TENN at 5 DIV. The two aggregates have been individually transduced to express GFP (left) and mCherry (right), allowing for identification and monitoring of aggregate-specific processes. **(H-K)** Confocal micrograph of the micro-TENN from **(G)**, showing axons projecting from each aggregate **(I, K)** and growing along each other **(J)**. Scale bars: 100 μm.

To advance micro-TENNs’ capabilities as an *in vitro* test-bed and/or to rebuild the damaged connectome, one of our design goals is to develop a computational platform that can be used to design and optimize micro-TENNs for specific performance goals. To be able to investigate neuronal growth, neurite extension, and the formation of synaptic connectivity at the distal ends, we need a simulation framework that can generate large-scale unidirectional and bidirectional axonal outgrowth with realistic neuron morphologies. The applications of this computational framework in micro-TENNs include: (i) study processes involved in outgrowth and structural integration in 3D microenvironments; (ii) aid in the design and optimization of functional characteristics and predict performance (e.g., output for a given input); (iii) simulate detailed neuron morphologies and anatomically-relevant neuronal-axonal networks to study connectome-level functional connectivity via spiking or compartmental network modeling. Combining the anatomical simulation results and the study of functional connectivity will increase our ability to understand and predict the neurophysiological characteristics and network-level activity in the micro-TENNs.

There are two major approaches to simulate neuronal development: construction algorithms and biologically-inspired growth processes [7]. Construction algorithms aim to reproduce the shape of real dendritic trees from distributions of shape parameters [8], [9]. However, this approach lacks the insight into any underlying biophysical mechanisms, such as the influences on morphological development caused by different neuronal types [10], a neuron’s intracellular environment and interaction with other neurons. Stochastic growth models, which provide a description of the growth process based on probabilistic growing events [11]–[14], is a popular approach under construction algorithms. Biologically-inspired growth processes are based on a description of the underlying biophysical mechanisms of the dendritic development [10], [15], [16]. The studies were conducted within various aspects of development, such as cell migration [17], neurite extension [18], growth cone steering [19], [20] and synapse formation [21].

In this paper, we present an ad-hoc growth model built upon the diffusion principle, which incorporates the stochastic process to reproduce the shape of micro-TENNs tissue. In that sense it belongs to the construction algorithms, however it does not rely on experimentally determined shape parameters. Our approach uses the 3D diffusion equation imposed with various rules for individual neuronal growth, such as the actions of neurite extension, branching, turning and aggregation/bundling. The concentration gradients guide the development of the axonal and dendritic neurites and describe the competition for resources between different growth tips of individual dendrites or axons.

## Methods

### Micro-TENN Fabrication and Experimental Measurements

Micro-TENNs were generated as previously described [4]. Briefly, agarose (3% w/v) was cast in a custom-designed acrylic mold to yield microcolumns with an outer diameter of 345 or 398 μm and inner diameter of 180 μm. Microcolumns were UV-sterilized and cut to a specified length before the lumen was filled with an ECM comprised of rat tail collagen 1 (1 mg/mL) and mouse laminin (1 mg/mL) adjusted to a pH of 7.2-7.4 (Reagent Proteins, San Diego, CA). To create the neuronal aggregates, embryonic day 18 (E18) cortical neurons were isolated from rodents and dissociated. The resultant single-cell suspensions were added to custom PDMS pyramidal wells and centrifuged at 200 x *g* for 5 minutes to force the cells into spheroidal aggregates. Following 24h incubation at 37°C/5% CO_2_, aggregates were seeded within the microcolumns to generate unidirectional (with one aggregate) and bidirectional (with one aggregate at each end) micro-TENNs. Micro-TENNs were then grown at 37°C/5% CO_2_ with half-media changes every 48 hours. To fluorescently label aggregates, adeno-associated virus 1 (AAV1) was sourced from the Penn Vector Core (Philadelphia, PA), packaged with the human synapsin 1 promoter and either green fluorescent protein (GFP) or the red fluorescent protein mCherry, and added to the pyramidal wells containing the aggregates (final titer: ~3×10^10^). Aggregates were kept at 37°C, 5% CO_2_ overnight before being seeded in micro-columns as described.

During the design and early development of the model, unidirectional and bidirectional micro-TENNs were generated with approximately 15-30E^3^ neurons per aggregate and lengths ranging from 2.0-9.0mm (n = 39), with growth rates analyzed as described [5]. To identify aggregate-specific axons over time, a set of 3.0mm-long, bidirectional “dual-color” micro-TENNs were simultaneously generated such that one aggregate expressed green fluorescent protein (GFP) while the opposing aggregate expressed mCherry (n = 6). Finally, for quantitative validation of the growth model, 2.0mm-long, unidirectional micro-TENNs were transduced to express GFP and generated with approximately 20E^3^ neurons per aggregate (n = 6) or 8.0E^3^ neurons per aggregate (n = 6) for characterization as described below. Micro-TENNs were imaged under phase contrast microscopy (magnification: 10x) at 1, 3, 5, 8, and 10 days in vitro (DIV) using a Nikon Eclipse Ti-S microscope paired with a QIClick camera and NIS Elements BR 4.13.00 (National Instruments). In addition to phase contrast microscopy, the bidirectional dual-color micro-TENNs were imaged at 1, 2, 3, 5, and 7 DIV using a Nikon A1RSI Laser Scanning confocal microscope paired with NIS Elements AR 4.50.00.

To quantify micro-TENN growth rates over time, the longest identifiable axons were measured from phase images at each DIV using ImageJ (National Institutes of Health, MD). Lengths were measured from the leading edge of the source aggregate (identified at 1 DIV) to the neurite tip, and growth was measured until axons from the aggregate either spanned the micro-TENN length (unidirectional) or began to grow along axons from the opposing aggregate (bidirectional). Growth rates were averaged at each timepoint to obtain a growth profile for unidirectional micro-TENNs with 20E^3^ and 8.0E^3^ neurons/aggregate. The peak growth rates for each group were compared using an unpaired t-test, with p < 0.05 set as the baseline for statistical significance.

To characterize axonal density with respect to cell count, phase images of unidirectional micro-TENNs with either 20E^3^ (n = 6) or 8.0E^3^ (n = 6) neurons/aggregate at 5 DIV were imported into ImageJ. 10-μm long rectangular regions of interest (ROIs) spanning the inner diameter (final ROI dimensions: 180 μm x 10 μm) were taken at 50% and 75% of the micro-TENN lengths. The axon density at these two locations was quantified as the percentage of the ROI populated by axons. Densities were averaged for the 20E^3^ and 8.0E^3^ groups and compared at each location via unpaired t-test with p < 0.05 as the baseline for significance. All data presented as mean ± s.e.m.

To characterize axon distribution, unidirectional micro-TENNs were fabricated and labeled with GFP (n = 5). At 10 DIV, micro-TENNs were gently drawn into a 22-gauge needle and vertically injected into a block of “brain phantom” agarose (0.6% w/v). Micro-TENNs were injected such that the aggregate was ventral with axon tracts projecting downward. Post-injection, micro-TENNs were imaged on a Nikon A1RMP+ multiphoton confocal microscope paired with NIS Elements AR 4.60.00 and a 16x immersion objective. Micro-TENNs were imaged with a 960-nm laser, with sequential 1.2μm-thick slices taken along the longitudinal axis (i.e. X-Y projections along the micro-TENN length). Post-imaging, the X-Y projections were used to generate a 3D reconstruction of the micro-TENN; cell bodies, axon bundles, and single axons were then manually identified via co-registration of the X-Y projections and 3D structure.

### Computational Model Development

The elongation and the growth direction of the neurites in the model is guided by concentration gradients. Each tip of each neuron is a diffusion source in free space. The bifurcation of the neurites is assumed to be a stochastic process, i.e. branching is associated with a time dependent probability function at each node. This framework aims to emulate the growth and bifurcation of micro-TENN neurons, however by using simple diffusion principles, it avoids the underlying biological complexity. All of the tips of the neurite tree are assumed to participate in the extension and branching process. Furthermore, extension and branching of each node are modeled as independent processes. This has computational advantages such as improved speed and ability to parallelize on a large scale.

The model uses continuous space/discrete time approach to allow freedom in the outgrowth direction and elongation. Space is bounded by the inner diameter of the hydrogel micro-column. The diameter and length of the tubular hydrogels, 180 μm and 2 mm respectively, are based on experiments previously performed by the Cullen Lab [1], [3]. In the micro-TENNs, axonal extension was measured approximately every two days; as such, the size of the fixed time interval of the model is 1% of this two-day interval (i.e. 28.8 minutes). In each time step, each individual axonal tip may (i) extend, (ii) bifurcate into two daughter branches and (iii) change growth direction. In the present implementation, the model uses fixed time steps with functions built upon the diffusion equation and concentration gradients for extension, turning, and branching. The model is developed with the condition that extension rate and turning direction depend on the concentration gradients at the terminal segment of each axon. The extension rate decreases exponentially to zero value [12] as the neurites stop growing due to the limitation of space and essential biochemical factors [14]. Branching probabilities are growing as a function of the simulation time.

#### Modeling Setup: Diffusion Equation and Concentration Gradient

Many stochastic models of neuronal activity are based on the theory of diffusion processes [22]. Several models have been developed to describe the growth of single neurons using the theory of one-dimensional stochastic diffusion [23]–[27].

In our bidirectional growth model, the tips of the neurons are diffusion sources in free space, assuming a constant isotropic diffusion coefficient. The governing equation is:

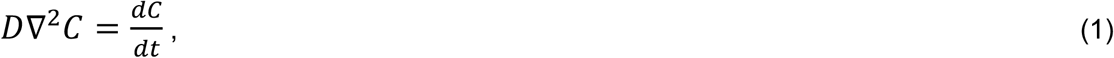

where *D* is the diffusion coefficient (*mm*^3^/*s*), *C* is the concentration (*mol*/*mm*^3^), and *t* is time (*s*). In Cartesian coordinates the partial differential equation becomes:

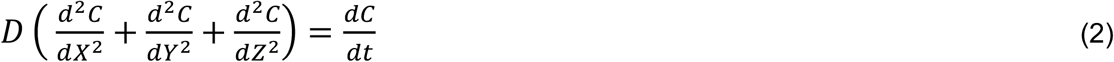

with boundary conditions:

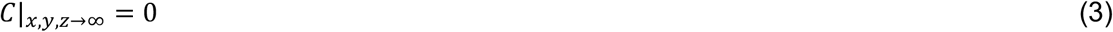

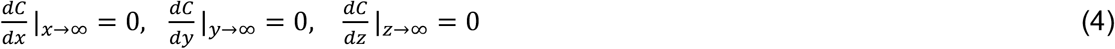

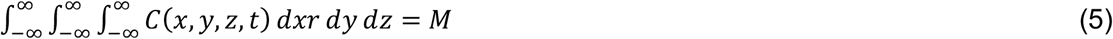

*M* is the initial amount of matter (*mol*). Without loss of generalization, we can choose *M* = 1 for convenience. The initial condition for a point source (*X*_0_, *Y*_0_, *Z*_0_) inside the shell is:

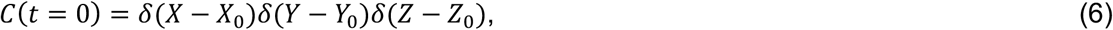

where δ(*X*) is Dirac’s delta function.

Thus, the general solution of the diffusion equation becomes:

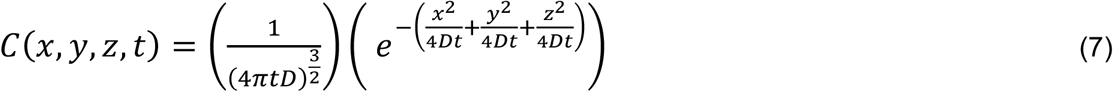

#### Direction of Neurite Outgrowth

The outgrowth of neurites is a complex process that is far from fully understood. In actual biological processes, the outgrowth direction of neurites depends on many intracellular and extracellular cues, which may cause large fluctuations in outgrowth directions [28], [29].

Our model is a Markov process: it assumes that the new outgrowth direction depends on the previous outgrowth direction and on the concentration gradients of the growth tips. For each growth tip, the concentration gradients are normalized to preserve the Markovian nature of the model.

The outgrowth direction is:

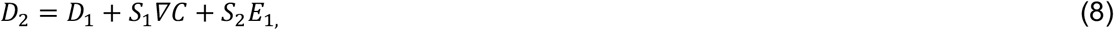

where *S*_1_ is the sensitivity to concentration gradients, *S*_2_ is the sensitivity to the direction perturbation, *D*_1_ is the previous direction vector, ∇*C* is the normalized concentration gradient, and *E*_1_ is the stochastic direction perturbation term.

Besides the gradients, a stochastic term *E*_1_ in Equation 8 is introduced to cause small fluctuations in the growth direction. Controlling this term in the simulation allows the control of the magnitude of deviation of the growth direction. Therefore, the component in the axial direction of the cylindrical tube has the largest value, while the components in the radial direction are relatively small.

#### Rate of Neurite Extension

The rate of extension of a neurite may vary considerably and is determined both by the external environment and by the internal state of the neurite [30]–[36]. In general, the extension rate decreases gradually with increasing distance from the soma [16]. In our model, the description of neurite extension rate follows the trend of experimental growth rate measured in unidirectional micro-TENNs. In each time step, the elongation of a single neurite is represented by the function,

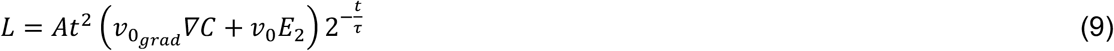

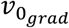 is the growth rate related to the gradients. *v* _0_ is the base extension rate and *E*_2_ is random process to cause fluctuation in *v*_0_. *t* is the simulation time, τ controls decreasing speed of the extension rate and ***A*** is a scaling factor.

#### Growth Tip Position

The coordinates at the next step of a growth tip are determined by the coordinates from the previous step, the outgrowth direction, and the extension rate. The new position in each time interval is given by

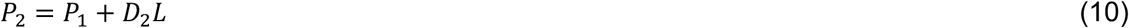

where *P*_1_ is the current tip position. However, this tip migration position cannot be accepted until it satisfies the coordinate restrictions of radial constraint and overlap avoidance.

#### Branching Probability, Rate of Branching, and Growth Rate After Branching

Neurite branching patterns are complex and show a large degree of variation in their shapes. Random branching on the segment indeed results in large and characteristic variations in the structures of the tree. As previous research has highlighted [12], [37], branching is assumed to occur exclusively at terminal nodes. Our model describes branching as a stochastic process. For each time step, for each of the terminal nodes in the growing tree, a branching probability *p*_q_ to form two new daughter nodes in a given time interval is assigned.

The probability of a branching event at each given terminal node *j* is given by:

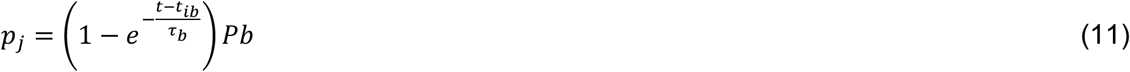

The time-dependent branching probability *p_j_* of a given terminal node *j* is dependent on several terms: the steady state branching probability *pb*(*pb* |_*t*_ = *∞*), simulation time *t*, the branching time step *t_ib_*, and a branching time constant *τ_b_*. The equation assumes that the branching probability of terminal nodes per time step remains constant for all tips. Branching probabilities are growing with the total simulation time. Such a function was necessary to match the shape of increasing number of dendritic terminal nodes during outgrowth of the micro-TENNs. The stochastic process of branching is also restricted by another random value *E*_3_; branching could only occur when both *p_j_* and *E*_3_ are greater than a certain value *B*. The value of *B* can be determined by the branching probability from experimental data. When a branching event takes place, two daughter terminal nodes are instantaneously added to the end of the existing terminal segment [38], which then becomes an intermediate segment.

The growth rate of the generated trees is closely related with segment outgrowth direction and extension. We only consider the extension distance in the axis direction as the growth distance. Thus, the growth rate is determined by the difference between the z components of the nodes. In the model, we force the growth cones to extend preferentially in the axis direction and the turning is relatively small. Therefore, the segment extension rate is the strongest factor to determine the growth rate. An estimate of υ_0_, and τ is obtained from the experimental growth rate. The optimization of the elongation parameters involves a comparison of the experimental and model segment extension rate.

With the current implementation, the branching probability increases with simulation time. The steady-state branching probability *pb* and time constant *τ_b_* are supposed to be extracted from experimental images. By controlling the values the values of *pb* and *τ_b_*, we have control over the morphology of the simulated neurites, since *pb* controls the branching density and *τ_b_*, controls how early in the process branching begins. In the extreme case of *pb*= 0, we can generate a morphology with no branching.

#### Radial Constraint

Experimentally, micro-TENNs were grown within miniature tubular hydrogels. In the simulation, the outgrowth process is also restricted within the tubular space (in this particular case, 180 micrometers in diameter [1], [3]), however the model allows different simulation radii to be employed. At each time step, the radial components of all the terminal nodes are tracked. If a radial component of a given node does not satisfy the tubular constraint, the node will be re-oriented to stick on the tubular wall.

#### Overlap Avoidance

The branches and extensions of neighboring neurites often target a shared or adjacent position. All the neurites are competing for space and avoiding overlap. Space competing is achieved by the concentration gradients. Overlapping is avoided by checking the distance from the new position to the surrounding existing segment tips. The model re-orients the growth direction when the extend position is sufficiently close to others.

#### Bundling and Helicity

The model accommodates fiber bundling. This is achieved through an attraction term *A*, i.e. the new position in each time interval becomes:

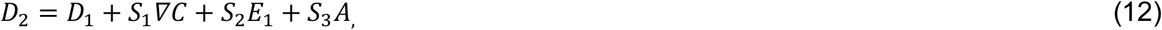

where *S*_3_ is a sensitivity to attraction (values between zero and one) that controls the bundle formation. In order to construct the attraction term *A*, we introduce an attraction radius of influence (*RI*). Every tip that falls within the *RI* is attracted to the centroids of all the tips that are within the *RI*. Larger values of *RI* lead to the formation of fewer bundles and vice versa. Thus, selection of different values of *RI* allows different morphologies with a different number of bundles. The model naturally provides an additional feature: tips belonging to a given bundle that fall outside of the *RI* at a given time step can form their own bundle, effectively allowing for a bundle to split. Such an effect is observed experimentally.

Helicity is another feature that was introduced to the model aiming to reproduce observed experimental micro-TENN morphologies. Very often, single axons and axonal bundles once reaching the inner wall of the micro-column, form a helix. Our model allows control over both the slope of the helix formed by an axon (or axonal bundle), as well as its helicity (handedness).

#### Parameter Optimization

Finding a best fit of the model-generated neuronal morphologies with an experimental data set requires an iterative comparison of experimental and model shape properties. In the search strategy, some parameters in the model are directly related to properties of the experimental data or images. For instance, the parameters 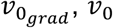, and *τ* predict the growth rate in axis direction. The branching process governed by parameters *τ_b_*, *P* and *B* fully determine the topological structure of the generated trees. These parameters are directly related to segment branching rate. The time step is selected to be ∆*t* = 0.02 days. Since υ_0_ is extracted from the experimental data, the selection of the value of the diffusion coefficient *D* is guided by the restriction:

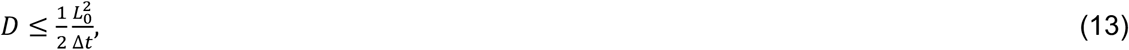

where *L*_0_= *L*(*t* = 0) is the initial extension.

Finally, the simulations were carried out on a laptop (Windows 10 Enterprise 64, Intel i7-7700HQ CPU @ 2.80GHz, 2801 Mhz, 4 Cores, 16GB DDR3 RAM). All code with is freely available at: https://github.com/PSUCompBio/GrowthModel.

## Results

### Examples of Micro-TENNs Morphologies

Variation of model parameters like sensitivity to attraction *S*_3_ and radius of influence *RI* allows us to generate different morphologies. *S*_3_ takes continuous values between zero and one, with *S*_3_= 0 corresponding to no bundle formation and *S*_3_= 1 corresponding to tight bundle formation. *RI* controls the number of bundles formed. **Figures 2, 3** and **4** demonstrate various unidirectional and bidirectional morphologies for different values of *RI* and *S*_3_ for micro-TENNs seeded with 3000, 6500 and 10000 cells.

**Figure 2.**
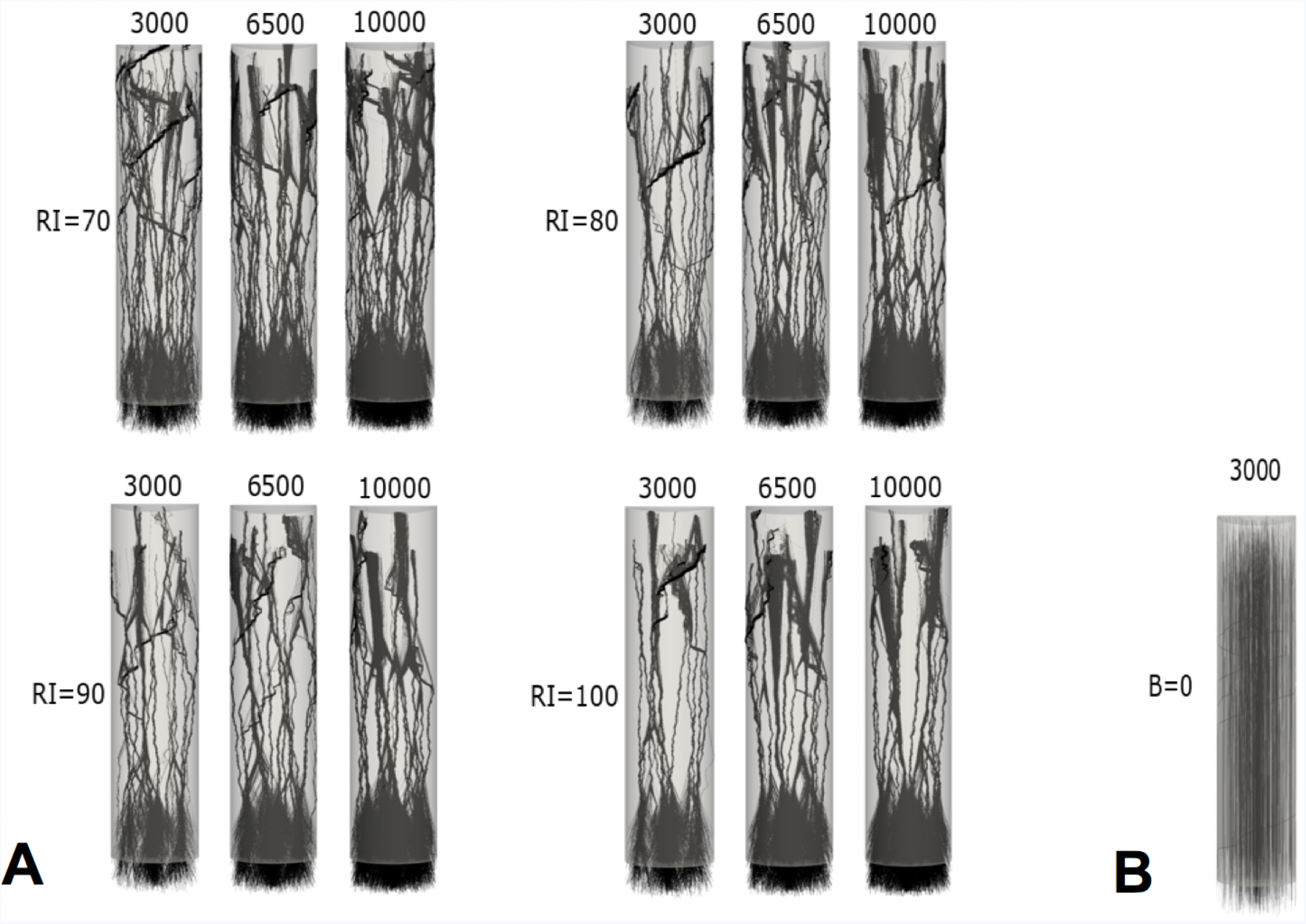
Example of unidirectional model-generated micro-TENN morphologies. (**A**) Twelve morphologies for micro-TENNs of length L=2000 *μm* and radius R=180 *μm*, number of seeded cells 3000, 6500 and 10000, and RI=70, 80, 90 and 100 *μm*, respectively. *S_3_*=1 causes tight bundle formation. (**B**) *S_3_*=0 guarantees no bundle formation.

**Figure 3.**
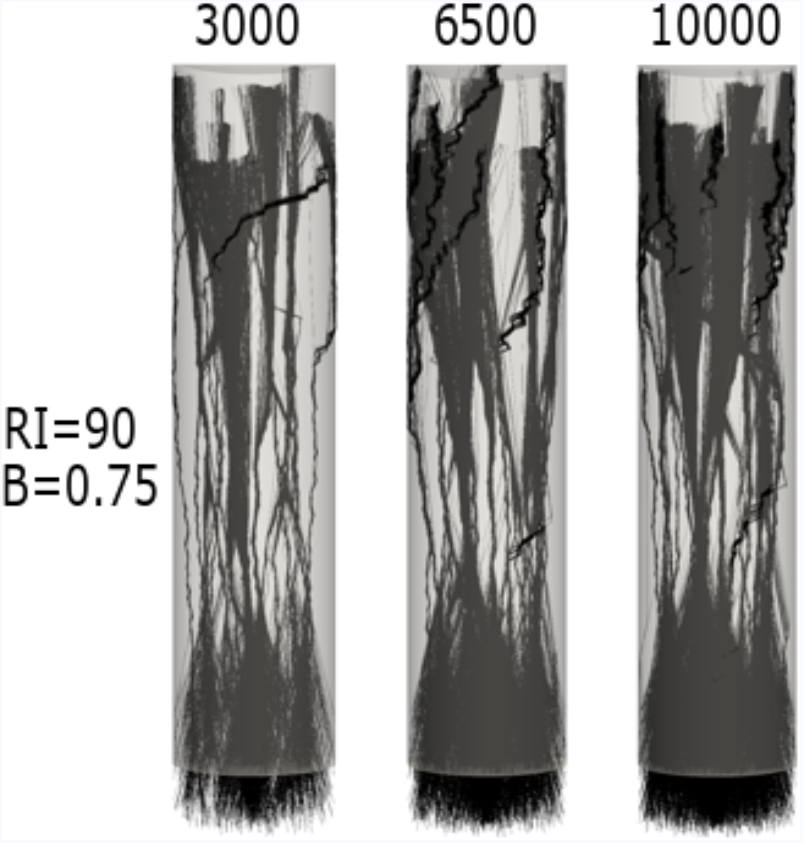
Example of unidirectional model-generated micro-TENNs. Micro-TENNs of length =2000 *μm* and radius R=180 *μm*, number of seeded cells 3000, 6500 and 10000, and RI= 90 *μm*. Here *S_3_*=0.75, bundles are not compact.

**Figure 4.**
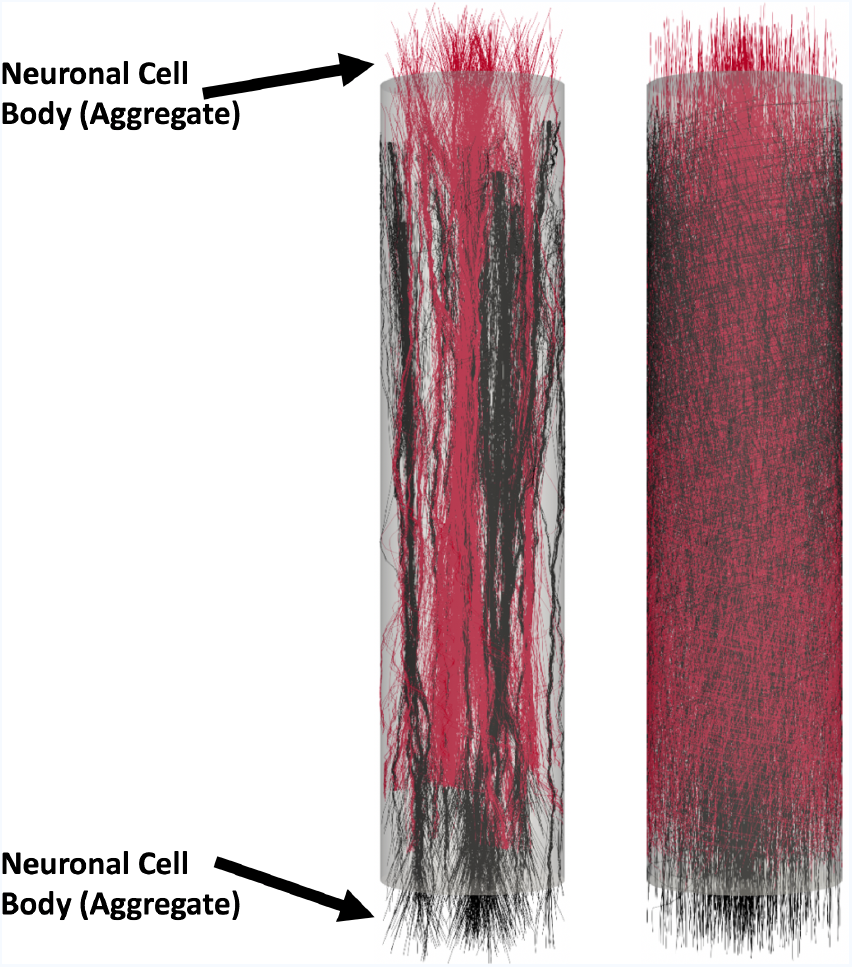
Example of bidirectional model-generated micro-TENNs. Analogously to the unidirectional case, the model can generate bidirectional morphologies. The micro-TENNs have length L=2000 *μm* and radius R=180 *μm*, the number of seeded cells is 3000. Here *S*_3_=1 and RI= 90 *μm* shows bidirectional axonal bundle formation (left), in the case *S*_3_=0 no bundles are formed (right). Neuronal cell bodies (aggregate) are labeled on each end but are hidden for clarity.

### Experimental Validation of Axonal Growth Rate

The model simulated the growth processes for unidirectional and bidirectional micro-TENNs when grown to 2000 *μm*. **Figure 5** shows images of unidirectional micro-TENNs of approximately 8000 and 2000 neurons, as well as measured and computed growth rate. The optimized parameters, *v*_0_ = 15, 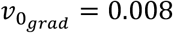 and *E*_2_ is a random uniform value between 0.8 and 1, provided an excellent fit with the experimental data. The result show that extension rate first increases, then decreases slowly with increasing distance from the soma. This is the trend for both the unidirectional and bidirectional growth rates.

**Figure 5.**
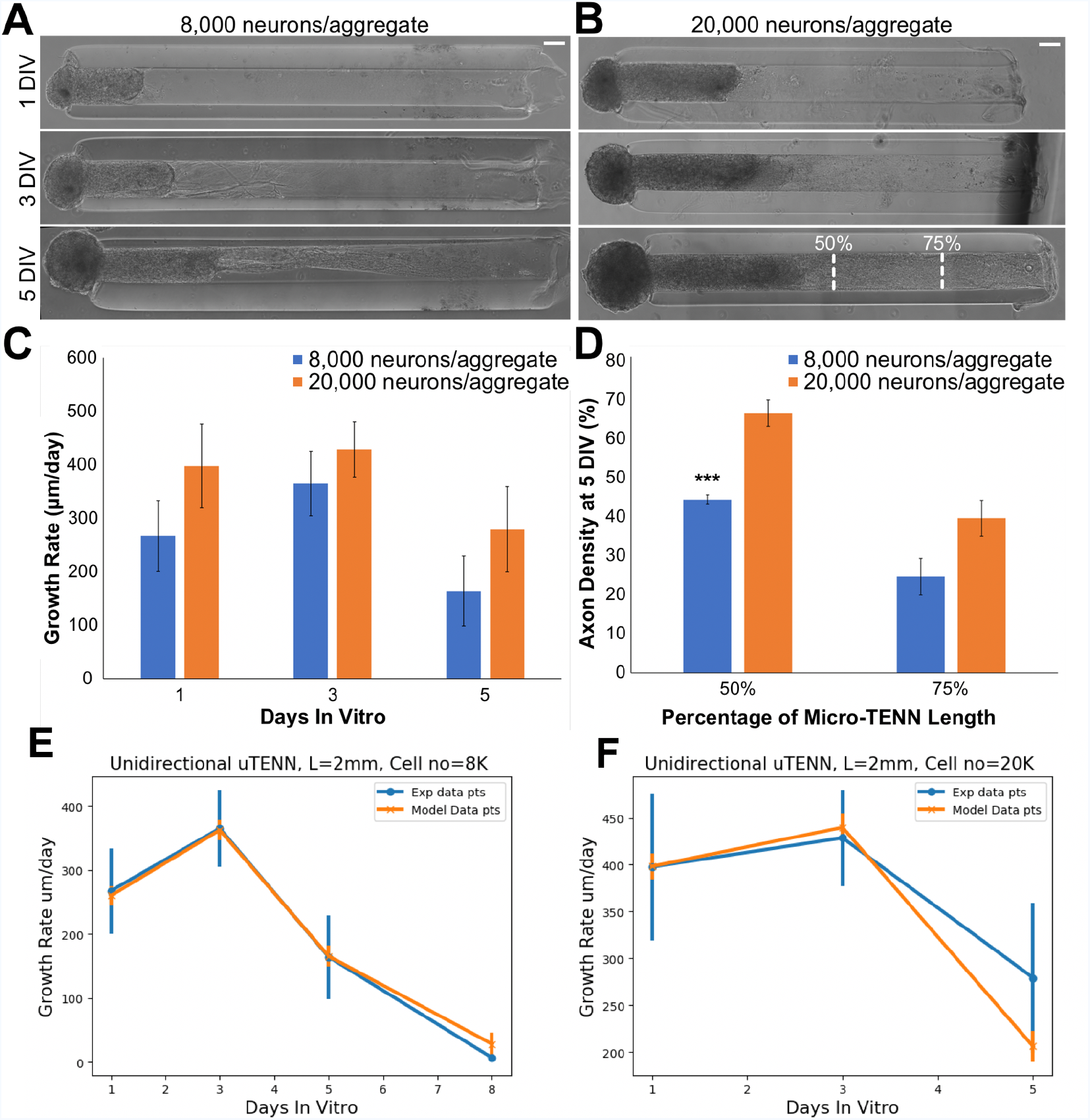
Validation of simulated growth rates *in vitro*. **(A)** Phase images of a unidirectional micro-TENN with approximately 8,000 neurons at 1, 3, and 5 DIV. **(B)** Phase images of a unidirectional micro-TENN with approximately 20,000 neurons at 1, 3, and 5 DIV. Both micro-TENN groups exhibited rapid axonal growth over the first few DIV. Scale bars: 100 μm. **(C)** Growth rates from both micro-TENN groups at 1, 3, and 5 DIV. Micro-TENNs with ~20,000 neurons/aggregate exhibited qualitatively faster growth rates than those with ~8,000 neurons/aggregate, although there were no statistically significant differences in growth rates. **(D)** Axon density at 5 DIV across the two groups at 50% and 75% along the micro-TENN length (as illustrated in (B)), quantified as the percentage of the microcolumn channel occupied by axons. Micro-TENNs with ~20,000 neurons/aggregate showed higher axon densities than those with ~8,000 neurons/aggregate, although this was only significant at 50% along the micro-TENN length (^***^ = p < 0.001). **(E)** Experimental versus model growth rate for unidirectional micro-TENN with 8,000 neurons. **(F)** Experimental versus model growth rate for unidirectional micro-TENN with 20,000 neurons.

**Figure 6A** shows a 3D reconstruction of a unidirectional micro-TENN at 10 DIV. Four corresponding slices were extracted for cross-sectional comparison to the computed results. **Figure 6 B-E** show each slice and provide arrows to distinguish between neuronal cell bodies, axon bundles, and single axons. In order to compare morphologies, we use two model generated X-Y projections along the Z-axis (**Figure 7**) that resulted from a micro-TENN simulation (inner radius 180 *μm*, length of 2mm, the number of seeded cells is 20,000). The model generates realistic morphology with single axons and axon bundles along the inner wall comparable to the experimentally reconstructed morphology in **Figure 7**.

**Figure 6.**
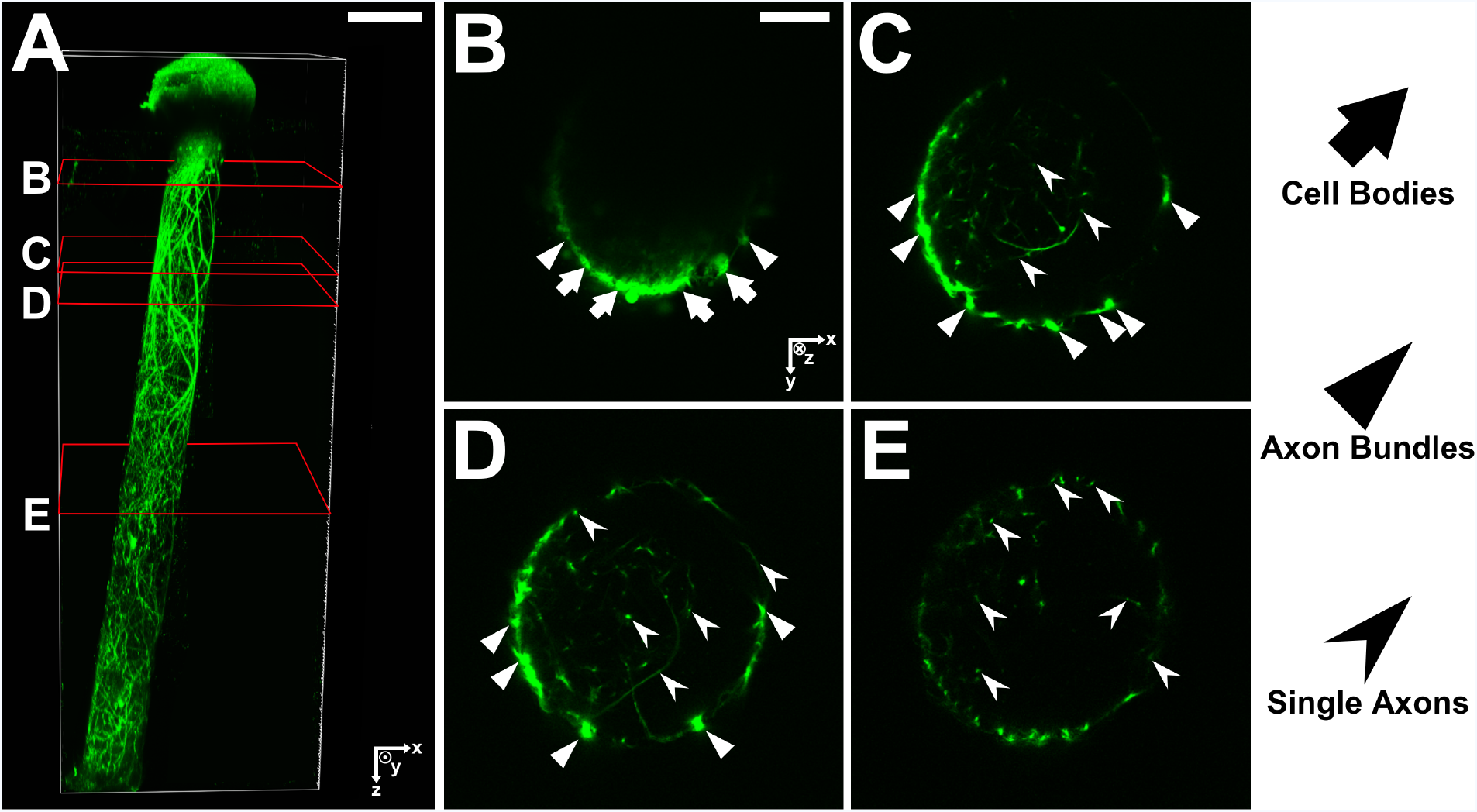
**(A)** 3D reconstruction of a unidirectional, GFP-positive micro-TENN at 10 DIV. **(B-E)** X-Y projections of the micro-TENN from (A) at the sections outlined in red. Orientation of the z-axis (positive) is into the page. Neuronal cell bodies (arrows) can be seen near the aggregate region in (B), from which axonal bundles (triangles) project and split into individual axons (caret). Scale bars: 200 μm (A); 50 μm (B).

**Figure 7.**
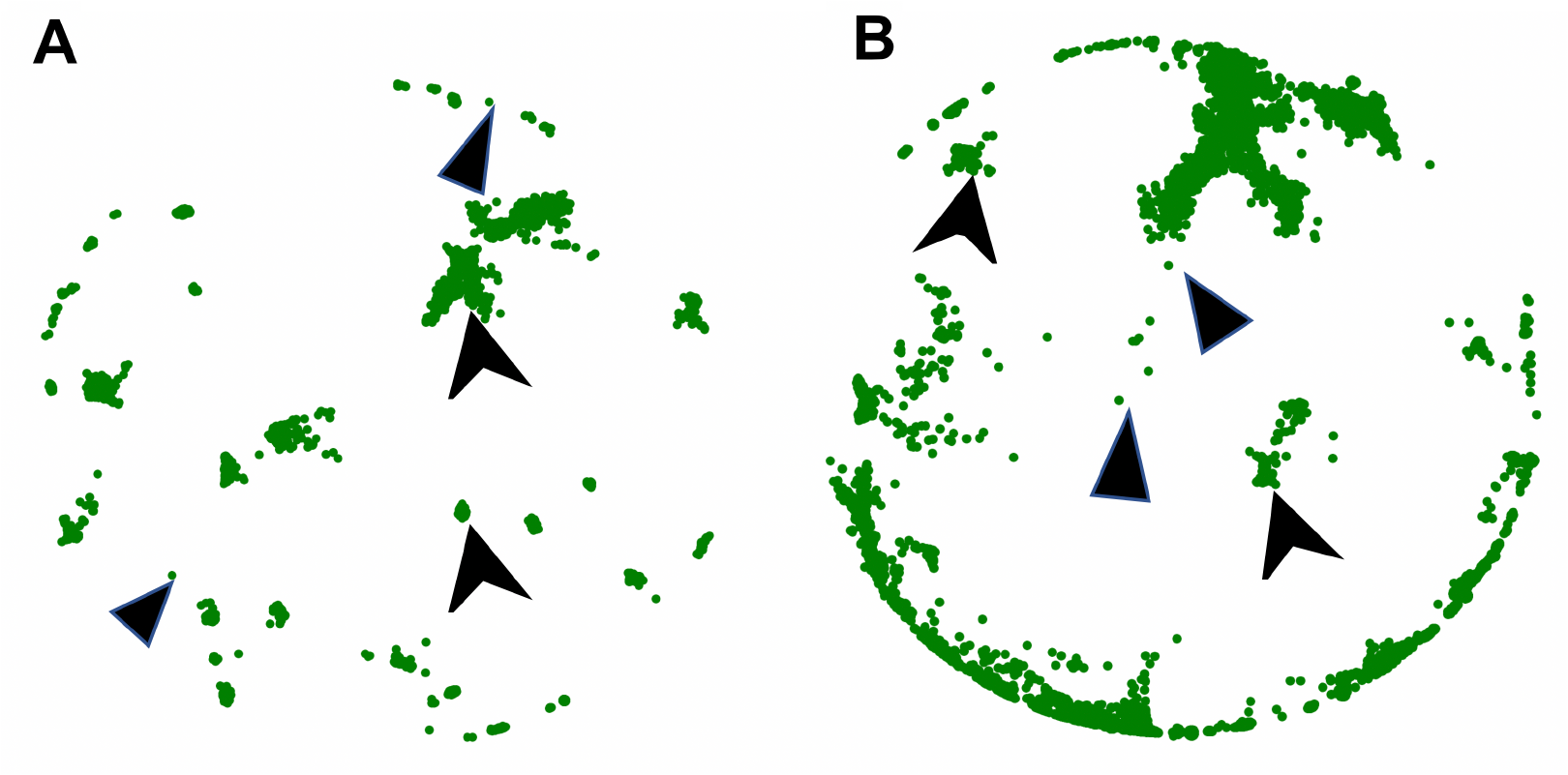
Model generated morphology of a unidirectional micro-TENN with inner diameter of 180 μm, length of 2mm and approximately 20,000 neurons. **(A)** X-Y projection at Z= 910 μm, model **(B)** X-Y projection at Z=1380 μm, model.

The computational algorithm, employed by our model is scalable as demonstrated on **Figure 8**, which shows the simulation runtime as a function of simulated number of cells (100, 1000 and 10000).

**Figure 8.**
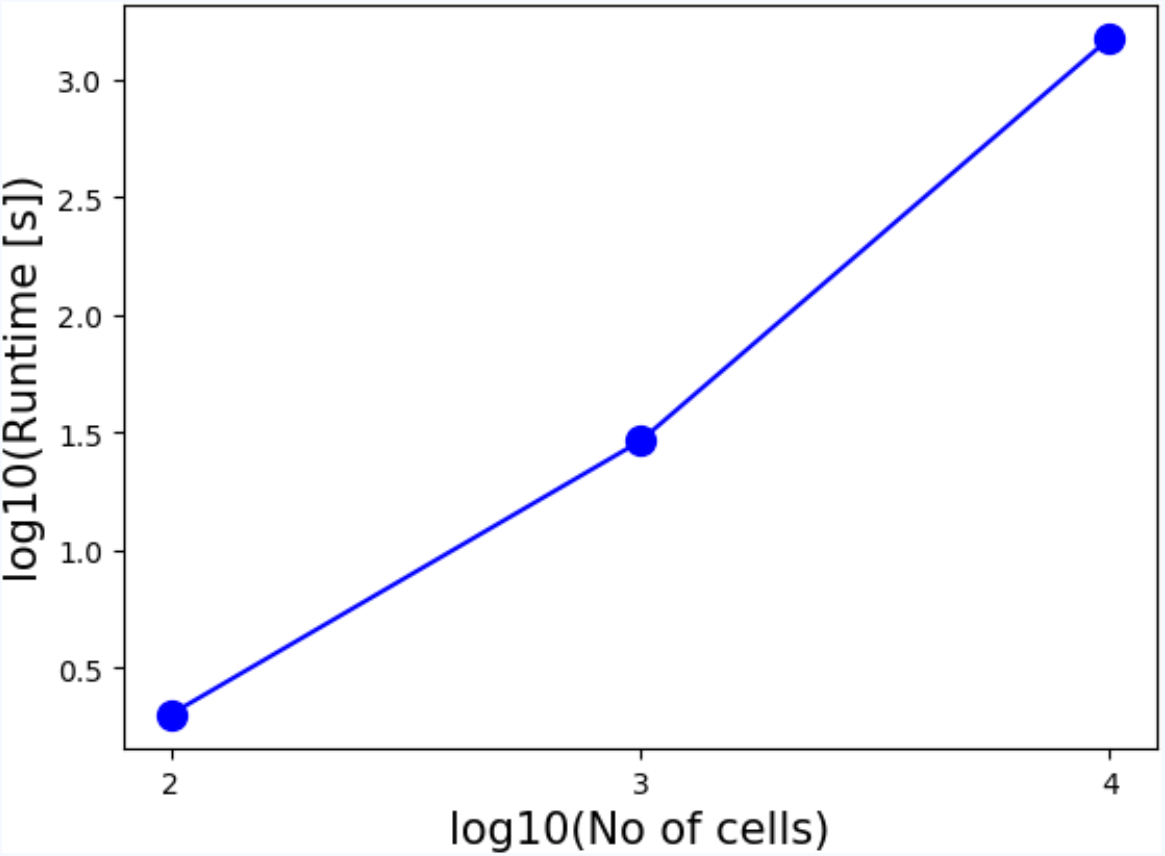
Scaling and computational speed. Simulations were performed for 100, 1000 and 10000 cells for the unidirectional micro-TENN with diameter 180 *μm* and length 2mm. The simulation runtime was 1.99 s, 28.93 s and 1500.45 s, respectively.

## Discussion

As a neural network model, micro-TENNs permit systematic interrogation of different contributors to neuronal growth and development in a three-dimensional, anatomically-relevant environment. Indeed, by providing precise control of the neuronal subtypes within the engineered aggregates, the extracellular matrix and milieu, as well as the potential presence of supporting glial cells, the micro-TENNs provide an ideal platform for the evaluation of interplay between intrinsic and extrinsic mechanisms of neuronal growth and neurite extension. For instance, the 3D biomaterial columnar encasement provides an unprecedented engineered environment to study the multi-faceted and often synergistic contributions of haptotactic [mediated by ECM (e.g., laminin, collagen) and cell-surface ligands (e.g., cadherins, L1)], chemotactic [mediated by growth factor gradients (e.g., nerve growth factor, glial derived neurotrophic factor) that can be attractive or repulsive], and mechanotactic [dictated by substrate geometry (e.g., curvature) and mechanical properties (e.g., stiffness)] on axonal outgrowth and pathfinding [39], [40]. To date, micro-TENNs have been generated with lengths ranging from 1-30 mm, and inner diameters as small as 160μm [4]. Moreover, the introduction of “actuator proteins” such as channelrhodopsin-2 (a light-sensitive ion channel for optically-induced neuronal stimulation) and/or activity markers such as the fluorescent calcium reporter GCaMP also provide a range of techniques to both modulate and monitor neuronal activity within the micro-TENN over time [5]. This controllability makes micro-TENNs an ideal testbed for eliciting and studying different neuronal phenomena under a range of experimental conditions, all within a three-dimensional architecture more similar to the native brain than traditional 2D cultures or randomly organized 3D cultures.

The existing models have been applied to study neuronal development *in vivo*, generally in the presence of molecular cues and under no specific geometric restrictions. Of note, those conditions differ from the growth conditions of micro-TENNs, in which gradients of external molecular cues are missing, the matrix is not neural tissue, and the growth space is a narrow tubular environment. Moreover, most of the existing models include complex growth mechanisms, leading to large computational cost. To compliment these previous efforts, there is a need for a computationally inexpensive model (due to the large population of neurons) that is capable of capturing the morphology of axonal growth within geometrical restrictions, including such important behaviors as neurite branching and axonal bundle formation/fasciculation.

Here we present a fast/computationally inexpensive ad-hoc stochastic process-based simulation framework for the generation of large-scale unidirectional and bidirectional neuronal networks with realistic neuronal-axonal morphologies. These simulations faithfully reproduce the shape of micro-TENNs, which are engineered microtissue networks formed by simultaneous axonal outgrowth of many neurons in a constrained (i.e. encapsulated) space.

The main advantage of the model is its conceptual simplicity. It is built on basic principles, yet it can generate various complex morphologies observed experimentally. Another major advantage is the computational speed. The solution of the diffusion equation for each tip is explicit and analytic, thus removing the necessity for a numerical solution for the concentration and the concentration gradients. This makes the model fast and computationally cheap, particularly for a large number of growing neurons. Also, each growth tip represents a separate process, allowing for parallelization and additional speed up of computational.

A limitation to the model is the introduction of parameters that cannot be extracted directly from experimental data. This can be due to the data resolution or simply to an inability of reliably quantifying certain experimental aspects. The parameter space they form has to be scanned for values that allow realistic neuronal morphologies. Another limitation of the model is the lack of chemical cues in the unidirectional case. While the model allows for additional attraction/repulsion and guidance terms to be introduced, employing direct extrapolation from *in vivo* growth models would be challenging.

The model is designed to capture some basic biological principles of neuronal development and axonal outgrowth *in vitro*: the competition for space and resources between growth tips, formation of bundles, chirality, the dependence of branching probability on the growth time, and the deceleration of the growth rate over time. The growth rate values in our model successfully reproduce the experimental data. Further development of the model could introduce additional guidance and attraction/repulsion molecular cues once such experimental information is available, thereby systematically adding complexity and the ability to capture synergistic and/or competing features of intrinsic and extrinsic growth parameters.

## Future Work

One major objective of building the Bidirectional Growth Model is to generate simulations of detailed neuron growth patterns to ultimately enable the study of functional connectivity that our research group has begun [41]. The neuronal growth patterns will be used to serve as the input of a spiking model to study the firing patterns within micro-TENNs and, following implantation, at the distal ends of micro-TENNs upon integration with the host brain neurons. In the output of the growth model, the framework can be used to extract detailed connectivity information.

The neuronal growth patterns provide the information for searching locations of synaptic connections and help to establish the spiking network simulation. In biological neuronal networks, synapses form where tissues are in sufficiently close proximity. According to experimental design, synapses occur close to the aggregate, which are in the 100 *μm* range from each end. Synaptic connectivity is estimated based on Euclidian distance (proximity criterion of 0.5 *μm*). **Figures 9A/2B** gives an example of the locations of these synaptic sites.

**Figure 9.**
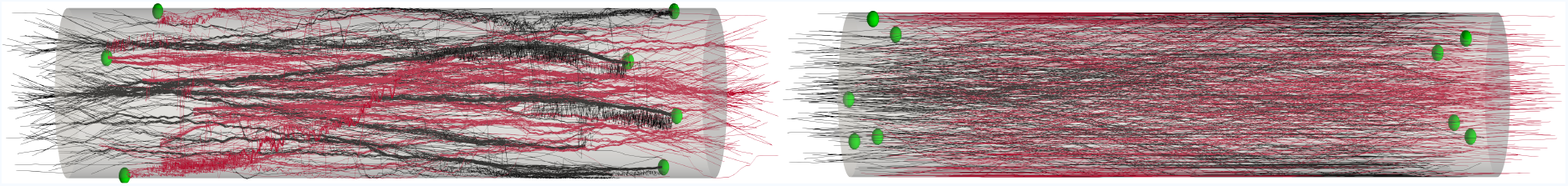
Examples of growth pattern and synapses. Illustration of synaptic formation close to the aggregate. The spheres indicate the synapses location. We consider that a synapse is formed when 2 fibers are closer than a threshold distance of 0.5*μm*. Synapses occur close to the aggregate (within 100 *μm*). The connectivity information is extracted for the spiking network simulation. (**A**) The output image of the model: synaptic formation in bidirectional micro-TENNs with axonal bundles. (**B**) The output image of the model: synaptic formation in bidirectional micro-TENNs with no axonal bundles.

## Conclusion

The neuronal and axonal growth structures obtained through this model provide a complete growth and connectivity pattern within a custom micro-tissue neural network. The model reproduces both the micro-TENN architecture and the axonal growth rate and distribution. This framework will enable further assessment of structural and functional connectivity, for instance an analysis of synaptic integration that happens close to the aggregate or even outside the micro-column. The extracted information of synaptic connectivity close to the aggregate and the synapse at distal end of micro-TENNs will be the a topic of a future functional connectivity study. We intend to build on this model in order to better understand the spiking network properties of micro-TENNs as so-called “living electrodes” for neuromodulation as well as anatomically-inspired constructs for white matter pathway reconstruction..

## Acknowledgements

Research reported in this publication was supported by the National Institutes of Health through the Brain Initiative [U01-NS094340 (Cullen & Kraft)] and the National Science Foundation through a Graduate Research Fellowship [DGE-1321851 (Adewole)]. Computations for this research were performed on the Pennsylvania State University’s Institute for CyberScience Advanced CyberInfrastructure (ICS-ACI).

